# *Botrytis cinerea* infection accelerates ripening and cell wall disassembly to promote disease in tomato fruit

**DOI:** 10.1101/2022.05.16.492169

**Authors:** Christian Silva, Jaclyn A. Adaskaveg, Saskia D. Mesquida-Pesci, Isabel B. Ortega-Salazar, Sivakumar Pattathil, Lisha Zhang, Michael G. Hahn, Jan A.L. van Kan, Dario Cantu, Ann L.T. Powell, Barbara Blanco-Ulate

## Abstract

Postharvest fungal pathogens benefit from the increased host susceptibility that occurs during fruit ripening. In unripe fruit, pathogens often remain quiescent and unable to cause disease until ripening begins, emerging at this point into destructive necrotrophic lifestyles that quickly result in fruit decay. Here, we demonstrate that one such pathogen, *Botrytis cinerea*, actively induces ripening processes to facilitate infections and promote disease. Assessments of ripening progression revealed that *B. cinerea* accelerated external coloration, ethylene production, and softening in unripe fruit, while mRNA sequencing of inoculated unripe fruit confirmed the corresponding upregulation of host genes involved in ripening processes, such as ethylene biosynthesis and cell wall degradation. Furthermore, an ELISA-based glycomics technique to assess fruit cell wall polysaccharides revealed remarkable similarities in the cell wall polysaccharide changes caused by both infections of unripe fruit and ripening of healthy fruit, particularly in the increased accessibility of pectic polysaccharides. Virulence and additional ripening assessment experiments with *B. cinerea* knockout mutants showed that induction of ripening is dependent on the ability to infect the host and break down pectin. The *B. cinerea* double knockout *Δbcpg1Δbcpg2* lacking two critical pectin degrading enzymes was found to be incapable of emerging from quiescence even long after the fruit had ripened at its own pace, suggesting that the failure to accelerate ripening severely inhibits fungal survival on unripe fruit. These findings demonstrate that active induction of ripening in unripe tomato fruit is an important infection strategy for *B. cinerea*.

**One-Sentence Summary:** Physiological, transcriptional, and glycomic evidence indicates *Botrytis cinerea* hastens tomato fruit ripening to facilitate infection by a mechanism dependent on the fungal enzymes Bcpg1 and Bcpg2.

## Introduction

Necrotrophic fungal pathogens often have broad host ranges and can cause disease in multiple tissues. Infections of fruit can display drastically different host-pathogen dynamics than those observed in vegetative tissues (Alkan and Fortes, 2015). Though both reproductive and vegetative tissues become more susceptible to necrotrophic pathogens during senescence (Häffner et al., 2015), in fruit, a dramatic increase in susceptibility is observed prior to senescence during ripening (Cantu et al., 2009; Prusky et al., 2013; Blanco-Ulate et al., 2016b). Because most fruit are economically valuable in their ripe state while vegetables are consumed prior to senescence, understanding ripening-associated susceptibility to disease is critical to reduce food losses and ensure high quality of fruit commodities.

Immature and unripe fruit are generally resistant to disease; however, some fungal pathogens can establish quiescent infections in these tissues (Prusky et al., 2013). The physiological nature of quiescence, including the level of pathogen colonization and activity, varies widely in different fruit pathosystems. For example, quiescence of the hemibiotrophic pathogen *Colletotrichum* in unripe fruit involves the development of melanized appressoria that penetrate and colonize a limited amount of fruit tissue (Guidarelli et al., 2011). In contrast, necrotrophic pathogens such as *Botrytis cinerea* are not known to produce such structures on fruit, yet they still survive in some capacity on the unripe fruit tissues before emergence from quiescence during fruit ripening (Adaskaveg et al., 2000; Petrasch et al., 2019a; Haile et al., 2020). Regardless of differences in quiescence, the onset and progression of fruit ripening trigger the pathogen to switch to an active necrotrophic lifestyle, resulting in rapid decay of fruit tissues.

Increased susceptibility to necrotrophs during fruit ripening is widespread across different fruit species and pathogens (Cantu et al., 2009; Alkan and Fortes, 2015; Petrasch et al., 2019b; Balsells-Llauradó et al., 2020). Fruit ripening is a complex suite of biophysical, physiological, transcriptional, and biochemical changes, and many of these are suspected to influence susceptibility to fungal pathogens (Blanco-Ulate et al., 2016b). Multiple such changes have been identified from the tomato-*B. cinerea* pathosystem, which has emerged as a model for fruit-necrotroph interactions (Cantu et al., 2008; Cantu et al., 2009; Blanco-Ulate et al., 2016b; Petrasch et al., 2019b; Silva et al., 2021). During ripening in tomato fruit, the pH of the apoplast decreases, providing a more favorable environment for the activity of virulence factors including proteases and cell wall degrading enzymes (Manteau et al., 2003). Ripening in tomato is also accompanied by a decline in antimicrobial compounds, such as α-tomatine (You and van Kan, 2021). However, one of the most significant contributors to susceptibility is the disassembly of the plant cell wall during fruit softening, given the importance of this structure as a physical barrier and source of plant defense signals (Cantu et al., 2008; Prusky et al., 2013; Blanco-Ulate et al., 2016a; Wang et al., 2022).

The cell walls of fruit generally have a higher proportion of pectins than hemicelluloses and cellulose, and fruit walls are usually more pectin-rich compared to walls of leaf and stem tissues (Brummell, 2006). During ripening, pectins are actively modified and degraded, hemicellulose and cellulose networks are loosened and broken down, and cell wall structural proteins are released or are no longer synthesized. In addition, the walls around cells in the pericarp and epidermis expand and become hydrated, leading to increased porosity of the cell wall structure and fruit softening (Brummell, 2006; Vicente et al., 2007). The relationship between endogenous host cell wall disassembly and host susceptibility is supported by the reduced susceptibility to *B. cinerea* observed in tomato mutant lines with suppressed or silenced expression of various cell wall degrading enzymes (CWDEs) and other related proteins including pectate lyase (*SlPL*; Silva et al., 2021) and the combination of polygalacturonase 2A (*S1PG2A*) and expansin 1 (*SlExp1*; Cantu et al., 2008).

The massive benefit of fruit ripening and host cell wall disassembly to fungal pathogens invites the possibility of active induction of ripening processes as an infection strategy to break quiescence. Some evidence indicates that *B. cinerea* may in fact do this in tomato: the pathogen induces host biosynthesis of the ripening-promoting hormone ethylene (Cantu et al., 2009; Silva et al., 2021) and increases expression of the host CWDEs *SlPG2A* and *SlExp1* in unripe tomato fruit (Cantu et al., 2008). However, a comprehensive examination of the extent to which *B. cinerea* induces ripening in unripe tomato fruit has not yet been performed. In this paper, we assess and compare the speed of various ripening processes, including color progression, ethylene production, and fruit softening, in mock-inoculated and *B. cinerea*-inoculated unripe tomato fruit. As a corollary, we sequenced mRNA in these tissues to detect the induction of metabolic pathways and genes associated with these ripening processes. To examine cell wall polysaccharide changes associated with fruit softening, we used an ELISA-based approach to compare changes in the fruit cell wall as a result of unripe fruit inoculation and ripening. Lastly, through virulence studies and additional ripening assessments, we determined that the combination of two *B. cinerea* genes, *Bcpg*1 *and Bcpg*2, is required for both ripening induction and emergence from quiescence in unripe tomato fruit.

## Results

### *Botrytis cinerea* infections accelerate ripening processes in unripe fruit

We hypothesized that inoculating unripe (mature green, MG) fruit with *Botrytis cinerea* would lead to accelerated ripening progression. To test this, we compared mock-inoculated and *B. cinerea*-inoculated MG fruit (*cv*. Ailsa Craig, AC) after several days post-inoculation (dpi). We first selected fruit at the MG stage based on size, color, and firmness, then divided these fruit randomly into two groups. Fruit from both groups were wounded six times. Then, fruit from the first group were inoculated with sterile water (i.e., mock-inoculated), while fruit from the second group were inoculated with a *B. cinerea* spore suspension We chose mock-inoculated MG fruit rather than healthy MG fruit to control for the effects of wounding. All fruit were then stored in high humidity and evaluated from 3 dpi to 6 dpi. These times of evaluation were chosen due to the fact that, up until 3 dpi, inoculated MG fruit remain resistant to disease and do not typically show signs of ripening (Silva et al., 2021).

Using non-destructive methods, we assessed the ripening rate in mock-inoculated and *B. cinerea-*inoculated MG fruit based on three characteristic physiological processes: external color progression, ethylene production, and loss of fruit firmness (i.e., softening). At 3 dpi, the first day of evaluation, mock-inoculated and *B. cinerea*-inoculated fruit were not significantly (*P* > 0.05) different for any of the ripening parameters evaluated, which confirmed that the fruit from both treatments were collected at an equivalent ripening stage (**Fig. 1**). *B. cinerea*-inoculated fruit, but not mock-inoculated, exhibited necrotic rings around the inoculation sites characteristic of response to *B. cinerea* in MG fruit (Cantu et al., 2008; Petrasch et al., 2019b). However, no advanced symptoms of fungal disease (e.g., water-soaked lesions or mycelial growth) were evident in any of the fruit at this initial time point.

**Figure 1:**
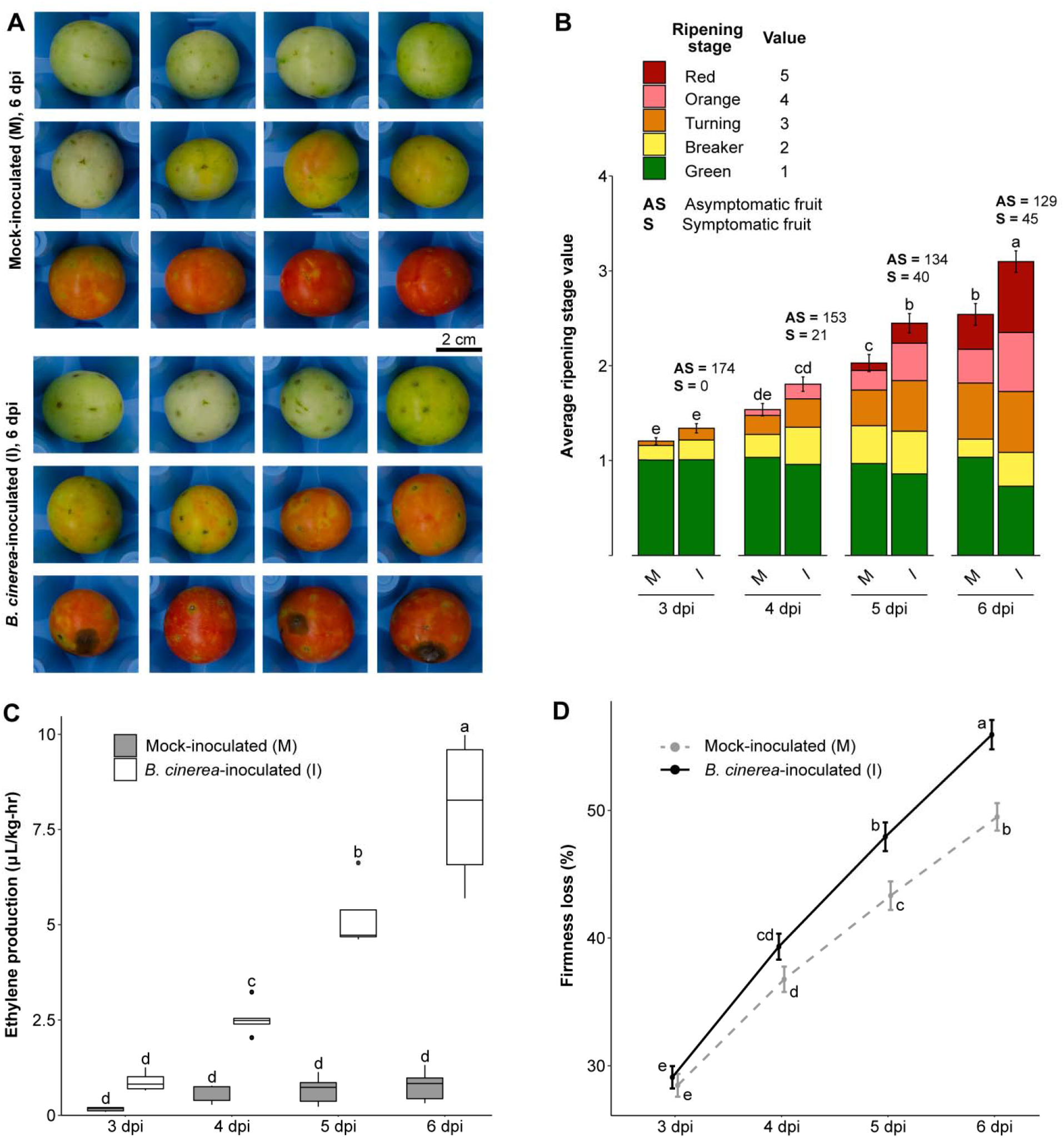
Acceleration of ripening in *B. cinerea*-inoculated unripe fruit. All panels contain data taken each day between 3 and 6 dpi. **(A)** Representative photos of mock-inoculated and *B. cinerea*-inoculated MG fruit at 6 dpi. (**B**) Average ripening stage value as assessed by color in mock-inoculated and *B. cinerea*-inoculated MG fruit (n = 174). Colored blocks within each column represent the proportion of fruit at the matching stage in the color key. (**C**) Production of ethylene in mock-inoculated and *B. cinerea*-inoculated MG fruit (n = 5). (**D**) Firmness loss in mock-inoculated and *B. cinerea*-inoculated MG fruit measured as a percentage of initial firmness at 0 dpi (n = 210 - 216). Letters in **B-D** indicate the statistical differences (*P* < 0.05) between each treatment across all dpi as calculated by ANOVA and Tukey’s HSD test. M = mock-inoculated, I = *B. cinerea-inoculated*.

After 3 dpi, ripening processes accelerated in *B. cinerea*-inoculated MG fruit. This coincided with a rise in the percentage of symptomatic inoculated fruit, reaching 26% at 6 dpi (**Fig. 1, Supplemental Table S1**). While fruit from both treatments experienced color progression based on their climacteric ripening behavior, *B. cinerea-*inoculated fruit turned red significantly faster (*P* < 0.05) than mock-inoculated fruit, which remained green longer (**Fig. 1**). For example, by 6 dpi, 44% of *B. cinerea-*inoculated fruit were at either the orange or red stage, compared to just 28% of mock-inoculated fruit. A rapid increase in ethylene production was observed only in the *B. cinerea-*inoculated fruit, where levels increased dramatically from 3 dpi onwards (*P* < 0.05); while ethylene levels remained constant in mock-inoculated fruit, indicating the normal climacteric ethylene burst had not yet occurred in most of these fruit (**Fig. 1**). Lastly, while both mock-inoculated and *B. cinerea*-inoculated fruit experienced a steady loss of firmness, fruit inoculated with the fungus lost firmness at a significantly (*P* < 0.05) faster rate, reaching an average of 56% loss compared to 49% in mock-inoculated fruit at 6 dpi (**Fig. 1**). Altogether, these results indicate that *B. cinerea* inoculations accelerated ripening processes, even when most of these fruits did not display any disease symptoms yet.

### *B. cinerea* infections induce premature expression of ripening-related genes in unripe fruit

We performed an RNAseq analysis to identify genes or pathways that could explain the accelerated ripening observed in the *B. cinerea*-inoculated MG fruit. We hypothesized that most physiological changes observed from 4 dpi onwards were preceded by a large transcriptional reprogramming in the fruit. Therefore, we proceeded to sequence mRNA from mock-inoculated, *B. cinerea*-inoculated, and healthy MG fruit (i.e., not wounded) after 3 dpi or 3 days post-harvest (dph). Compared to healthy fruit, we expected to see induction of ripening-related transcriptional activity by *B. cinerea* inoculation, but not mock inoculation. We also incorporated two other existing transcriptomic datasets: (i) mock-inoculated, *B. cinerea*-inoculated, and healthy samples created identically to the 3 dpi samples but sequenced at 1 dpi (Silva et al., 2021) to account for the possibility that genes were triggered earlier during the inoculation, and (ii) publicly available samples of healthy fruit at five developmental stages from MG to red ripe (RR) from the fruitENCODE database (Lü et al., 2018) to capture ripening-related genes.

When compared to healthy MG fruit, 5,512 genes were found to be differentially expressed (*P_adj_* < 0.05) as a result of *B. cinerea* inoculation (MG I / MG H) at either 1 or 3 dpi. In contrast, a much smaller number of genes (582) were differentially expressed due to mock inoculation (MG M / MG H), and most of these (482 or 82.8%) had the same expression pattern as they had in *B. cinerea*-inoculated fruit (**Table 1**). These results indicate that *B. cinerea* inoculation, but not mock inoculation, has a substantial and targeted impact on gene expression in MG fruit, and that wounding responses in MG fruit represent a small subset of fungal inoculation responses. From the fruitENCODE data, a total of 10,795 genes were found to be differentially expressed at one or more of the four ripening stages when compared to MG. These ripening genes were then used to determine if *B. cinerea* inoculation could induce similar transcriptional changes in MG fruit. A table of the differential expression results, including relevant gene annotations, can be found in **Supplemental Table S2**.

**Table 1:** Differentially expressed genes as a result of mock inoculation, *B. cinerea* inoculation of unripe fruit, and healthy fruit ripening. *Total values in these cells include genes with mixed expression patterns (i.e., upregulated in one subcategory and down in another subcategory) within that category. B = Breaker, B + 5 = Breaker + 5 days, B + 7 = Breaker + 7 days.

| Category | Subcategory | Upregulated | Downregulated | Total |
| --- | --- | --- | --- | --- |
| **Mock Inoculation**<br>(MG M / MG H) | 1 dpi | 499 | 31 | 530 |
|  | 3 dpi | 139 | 24 | 163 |
|  | Total | 533 | 49 | 582* |
| ***B. cinerea* Inoculation**<br>(MG I / MG H) | 1 dpi | 2,823 | 1,734 | 4,557 |
|  | 3 dpi | 1,934 | 1,004 | 2,938 |
|  | Total | 3,249 | 2,252 | 5,512* |
| **Ripening** | B / MG | 2,341 | 2,761 | 5,102 |
|  | B + 5 / MG | 2,292 | 3,277 | 5,569 |
|  | B + 7 / MG | 3,543 | 4,491 | 8,034 |
|  | RR / MG | 3,383 | 4,644 | 8,027 |
|  | Total | 4,617 | 5,377 | 10,795* |

We first focused on the expression of genes with functional annotations belonging to three different categories: carotenoid biosynthesis, ethylene biosynthesis, and cell wall degrading enzymes (CWDEs), due to their link to the accelerated ripening processes that we demonstrated above (**Fig. 1**). We were particularly interested in looking at transcriptional changes caused by *B. cinerea* inoculation, fruit ripening, or both. Several genes in the lycopene biosynthesis pathway were significantly (*P_adj_* < 0.05) upregulated during ripening, including *phytoene synthase 1* (*SlPSY1, Solyc03g031860*), *zeta-carotene isomerase* (*SlZ-ISO, Solyc12g098710*), *zeta-carotene desaturase* (*SlZDS, Solyc01g097810*), and *carotene isomerase* (*SlCrtISO, Solyc03g007960*; **Supplemental Table S2**). Curiously, none of these genes appeared to be significantly up-regulated as a result of *B. cinerea* or mock inoculation at 1 or 3 dpi, though baseline expression of *SlPSY1* in MG fruit at 3 dpi was relatively high (average normalized read count = 46,519) in the MG fruit regardless of the treatment. The transcriptional induction of genes involved in color progression may not be evident in our RNAseq data due to indirect effects on the metabolic flux through manipulation of a neighboring pathway, non-transcriptional regulation, or simply transcriptional induction at a different time point than those evaluated.

*B. cinerea* inoculation resulted in a clear upregulation of ethylene biosynthesis genes (**Fig. 2**). Interestingly, while *B. cinerea* sometimes upregulated paralogs involved in ripening (*SlACS4, Solyc05g050010* and *SlACO1, Solyc07g049530*), it often upregulated additional ones that are not normally associated with ripening or are not involved in System 2 ethylene production (*SlACS8*, *Solyc03g043890*; *SlACO2*, *Solyc12g005940*; and *SlACO3, Solyc07g049550*). As ethylene is a strong promoter of fruit ripening in tomato, induced ethylene biosynthesis in *B. cinerea*-inoculated MG fruit may lead to activation of downstream ripening genes, thus accelerating the ripening process even further.

**Figure 2:**
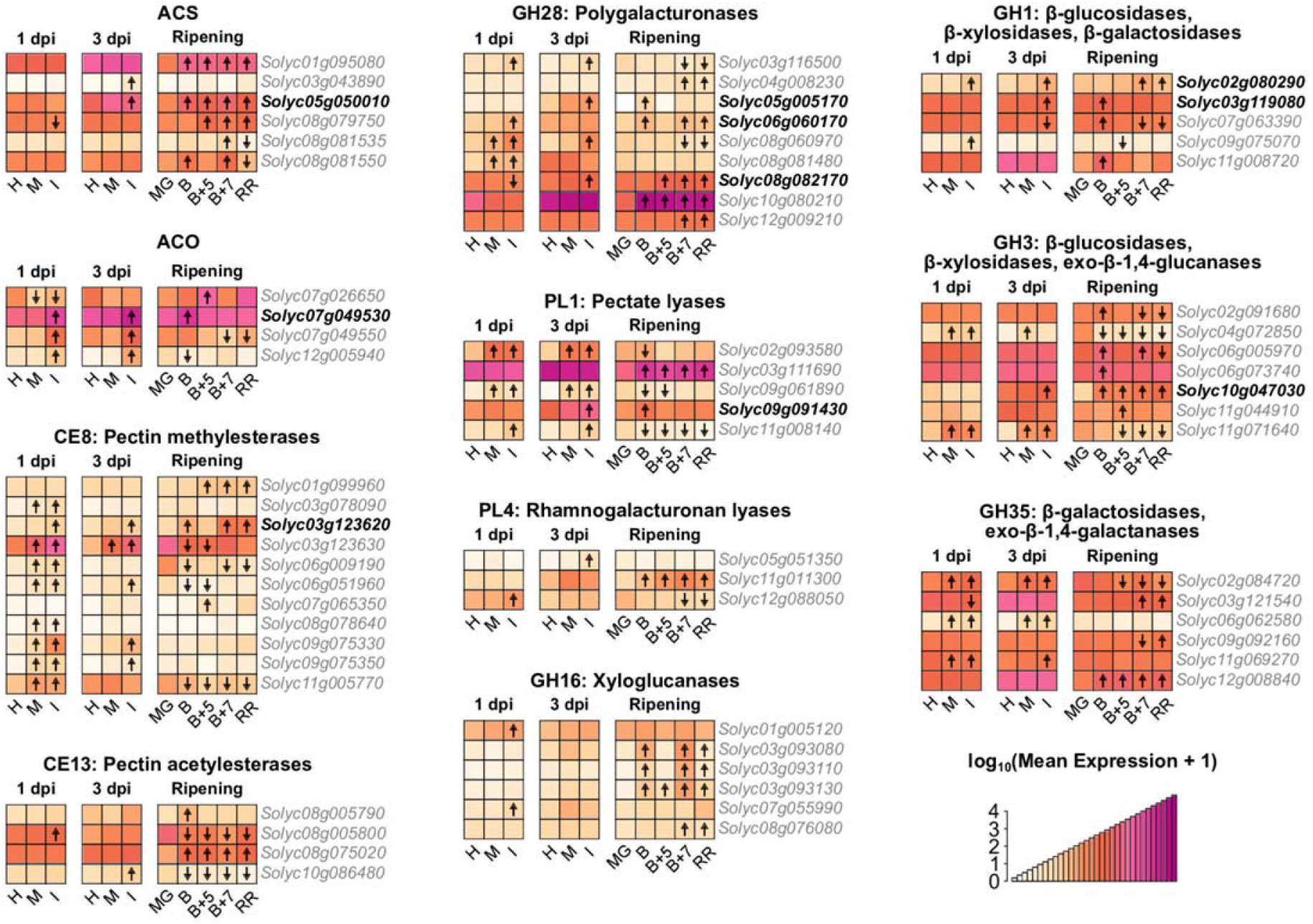
Expression patterns of ethylene biosynthesis and cell wall degrading enzyme genes in *B. cinerea*-inoculated unripe fruit and healthy fruit ripening. Heatmaps of normalized expression values in healthy, mock-inoculated, and *B. cinerea*-inoculated MG fruit at 1 and 3 dpi, as well as healthy ripening obtained from the fruitENCODE (Lü et al., 2018). Normalized expression values have undergone a log_10_(mean expression + 1) transformation. Arrows within heatmap tiles indicate statistically significant (*P_adj_* < 0.05) upregulated or downregulated genes when compared to expression values in healthy MG fruit at that timepoint. Boldtype font indicates genes that were upregulated by both *B. cinerea* inoculation and ripening. H = healthy, M = mock-inoculated, I = inoculated, B = Breaker, B + 5 = Breaker + 5 days, B + 7 = Breaker + 7 days.

Likewise, CWDE genes from nine families known to be involved in fruit softening were induced by *B. cinerea*. A total of 30 genes across these families were found to be upregulated during ripening, with eight of these genes also being upregulated by *B. cinerea* inoculation. However, *B. cinerea* inoculation induced an additional 26 CWDEs beyond the eight ripening-related ones, demonstrating substantial recruitment of host CWDEs by the pathogen after MG inoculation. These included nine pectin methylesterases (PMEs), many of which appeared to be also induced by mock inoculation at 1 dpi, though by 3 dpi upregulation was sustained by only *B. cinerea* inoculation for all but one gene. Also upregulated by *B. cinerea* were six polygalacturonases (PGs) and three pectate lyases (PLs), enzymes responsible for pectin backbone depolymerization. *B. cinerea* only weakly upregulated two xyloglucanases at 1 dpi. Additional glycosyl hydrolases with mixed activity on pectin, hemicellulose, cellulose, and other sugars (GH1, GH3, and GH35) were also found to be upregulated during ripening and *B. cinerea* infection. Altogether, these results suggest that *B. cinerea* infections lead to the activation of host CWDE expression, particularly pectin-related enzymes, which then facilitate the disassembly of the fruit cell walls.

To confirm that *B. cinerea* inoculation results in meaningful ripening gene expression changes beyond 3 dpi, we selected seven genes in the carotenoid biosynthesis, ethylene production, and cell wall degradation pathways for further assessment via quantitative PCR (qPCR). All genes exhibited significantly (*P* < 0.05) greater expression in *B. cinerea*-inoculated MG fruit compared to mock-inoculated MG fruit for at least two of the later timepoints evaluated (3 dpi, 4 dpi, 5 dpi, or 6 dpi) (**Supplemental Figure 1**). Notably, this included the carotenoid biosynthesis genes *SlPSY1* and *SlZDS*, which were not significantly induced in *B. cinerea*-inoculated fruit at 3 dpi in our RNASeq analysis (**Fig. 2**). The qPCR data confirmed the prominent expression of ethylene biosynthesis genes *SlACO1, SlACO3*, and *SlACS8* in *B. cinerea*-inoculated fruit, in contrast to the lower expression of these genes in mock-inoculated fruit even at 6 dpi, underscoring the relative lack of ethylene biosynthesis in these fruit (**Fig. 1**). Furthermore, we detected that the ripening-related CWDEs *SlPG2a* (*Solyc10g080210*) and *SlPL* (*Solyc03g111690*) were upregulated by *B. cinerea* inoculation after 3 dpi.

Beyond the carotenoid biosynthesis, ethylene biosynthesis, and CWDE categories, we were interested in the overall overlap between genes induced by *B. cinerea* inoculation and the ripening-related genes. A total of 629 genes were commonly upregulated during MG inoculation (1 and/or 3 dpi) and ripening, and 1,031 genes were downregulated in these comparisons. To identify prevalent functions of these genes, we performed enrichment analyses (*P_adj_* < 0.05) of KEGG pathway annotations (**Table 2**). Among the commonly upregulated genes, the most significantly enriched pathways were “plant-pathogen interaction” (sly04626) and “proteasome” (sly03050). The sly04626 genes were found to mostly consist of various calmodulins and calcium-dependent protein kinases. Additionally, the “alpha-linolenic acid metabolism” (sly00592) pathway was enriched, and the corresponding genes were found to be those responsible for the biosynthesis of jasmonic acid, a hormone that, similarly to ethylene, positively regulates ripening and pathogen responses. Commonly downregulated genes revealed an abundance of various photosynthesis-related pathways. The decreased photosynthetic capacity as a result of ripening fruit is well-known, and the occurrence of this as the result of *B. cinerea* inoculation can help explain the accelerated color progression in the inoculated MG fruit. Ultimately this overlap between MG inoculation responsive genes and ripening-related genes indicates that *B. cinerea* may activate multiple ripening processes.

**Table 2:** Pathway enrichment among genes commonly upregulated or downregulated by both *B. cinerea* inoculation of unripe fruit and ripening. Functional pathways were defined using the KEGG database, KEGG codes are given within parentheses. Only significantly enriched pathways (*P_adj_* < 0.05) are shown.

| Category | KEGG Pathway | Number of Genes | $P_{adj}$ |
| --- | --- | --- | --- |
| Upregulated | Plant-pathogen interaction (sly04626) | 18 | $2.9 \times 10^{-7}$ |
| | Proteasome (sly03050) | 10 | $2.0 \times 10^{-6}$ |
| | Glutathione metabolism (sly00480) | 10 | $2.7 \times 10^{-3}$ |
| | alpha-Linolenic acid metabolism (sly00592) | 6 | $2.0 \times 10^{-2}$ |
| | Citrate cycle (sly00020) | 6 | $3.8 \times 10^{-2}$ |
| | Sulfur metabolism (sly00920) | 5 | $4.7 \times 10^{-2}$ |
| Downregulated | Photosynthesis (sly00195) | 30 | $7.5 \times 10^{-21}$ |
| | Carbon fixation (sly00710) | 25 | $5.8 \times 10^{-18}$ |
| | Photosynthesis – antenna proteins (sly00196) | 16 | $9.7 \times 10^{-12}$ |
| | Pentose phosphate pathway (sly00030) | 13 | $4.4 \times 10^{-7}$ |
| | Glyoxylate and dicarboxylate metabolism (sly00630) | 12 | $1.2 \times 10^{-4}$ |
| | Fructose and mannose metabolism (sly00051) | 11 | $7.1 \times 10^{-4}$ |
| | Porphyrin and chlorophyll metabolism (sly00860) | 9 | $1.3 \times 10^{-3}$ |
| | Glycolysis / gluconeogenesis (sly00010) | 13 | $1.0 \times 10^{-2}$ |

### Healthy ripening and unripe fruit inoculation with *B. cinerea* result in similar changes to cell wall polysaccharide composition

Of the three ripening processes analyzed above, induction of cell wall degradation is likely to have the largest impact on the disease outcome by facilitating fungal colonization. We profiled the cell wall glycome of fruit to obtain a deeper understanding of the similarities between the cell wall changes induced by MG fruit inoculation and those that occur during normal ripening. For our comparisons, we selected three types of fruit: (i) *B. cinerea*-inoculated MG fruit at 3 dpi, (ii) healthy MG fruit at 3 dph, and (iii) healthy RR fruit at 3 dph. Because mock-inoculated fruit showed very limited induction of CWDE expression, most of which overlapped with the CWDEs induced by *B. cinerea*, we did not include them in these analyses. As with the transcriptomic data, we chose 3 dpi or 3 dph as our assessment time point because it is the last day before symptoms of the disease appear in MG inoculated fruit.

From the total cell wall material, we generated four different soluble fractions each differing in their polysaccharide composition. The water-soluble fraction (WSF) contained small molecules and pectin polysaccharides that are soluble in un-buffered water. The CDTA-soluble fraction (CSF) included calcium-bound pectins. The Na_2_CO_3_-soluble fraction (NSF) was composed of pectins linked to the cell wall matrix via covalent ester linkages. The KOH-soluble fraction (KSF) was enriched for hemicelluloses (xyloglucans and xylans). All fractions were subjected to glycome profiling to detect diverse epitopes present in pectin, hemicelluloses, or mixed polysaccharide substrates (**Fig. 3**).

**Figure 3:**
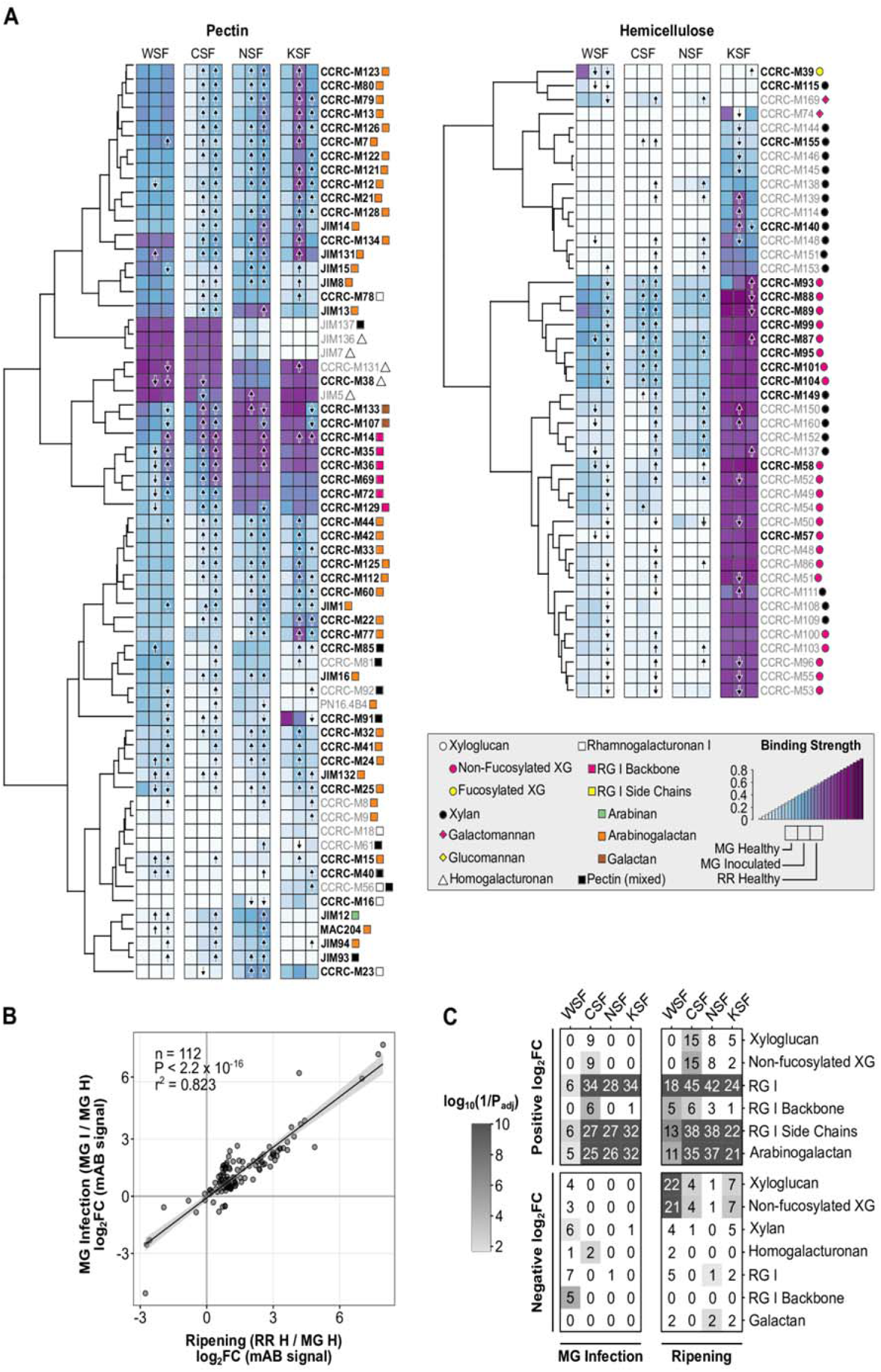
Glycomics profiling of *B. cinerea*-inoculated unripe fruit. (**A**) Heatmaps of binding strength of polysaccharide-binding mABs) in healthy MG, *B. cinerea*-inoculated MG fruit, and healthy RR at 3 dpi or dph. mAB codes are given to the right of each heatmap row, with the recognized classes of cell wall polysaccharides indicated by colored shapes according to the given key. mAB listed in boldtype are those included in the scatterplot in panel **B**. Arrows within heatmap tiles indicate statistically significant (*P_adj_* < 0.05) increasing or decreasing antibody strength when compared via t-test to values in healthy MG fruit (n = 6). (**B**) Scatterplot and linear regression model of log_2_ fold change (log_2_FC) values of mAB signals in MG inoculation (*B.cinerea*-inoculated MG / healthy MG) and ripening (healthy RR / healthy MG) comparisons. (**C**) Enrichment of polysaccharide classes with statistically significant positive or negative log2FCs in each cell wall fraction for the ripening and MG inoculation comparisons. Numbers within each tile indicate the number of mABs with a statistically significant log_2_FC in that respective fraction and polysaccharide class. WSF = water-soluble fraction, CSF = CDTA-soluble fraction, NSF = Na_2_CO_3_-soluble fraction, KSF = KOH-soluble fraction.

A total of 112 of 144 assessed monoclonal antibodies (mABs) assayed across the four different fractions demonstrated significant (*P* < 0.05) log_2_ fold changes for both MG fruit inoculation (MG I/MG H) and ripening (RR H/ MG H; **Fig. 3**). These log_2_ fold changes correlated strongly between comparisons (adjusted *r*^2^ = 0.82), suggesting a high degree of similarity in cell wall polysaccharide changes brought on by *B. cinerea* inoculation in MG fruit and fruit ripening. Nearly all (99/112) of these mABs showed increased binding (i.e., positive log2 fold changes) in both comparisons, indicating that MG fruit inoculation and ripening increase access to pectin polymers reflective of cell wall disassembly.

We performed enrichment analyses (*P_adj_* < 0.05) to identify overrepresented polysaccharide classes among the mABs with significant log2 fold changes in each fraction (**Fig. 3**). Enrichment patterns were remarkably similar between MG fruit inoculation (MG I / MG H) and ripening (RR H / MG H). In particular, multiple mABs associated with the rhamnogalacturonan (RG) I backbone and arabinogalactans experienced increased binding strength in the CSF, NSF, and KSF fractions in both comparisons. In contrast, changes in binding strength of hemicellulose-specific mABs (e.g., targeting non-fucosylated xyloglucans) were largely restricted to the ripening process, consistent with the relative lack of hemicellulose-specific CWDEs activated during infection (**Fig. 2**).

To test if the ripening-like polysaccharide changes due to *B. cinerea* inoculation were specific to MG fruit, we performed the same glycomics analyses with inoculated RR fruit (RR I / RR H; **Supplemental Figure 2**). Unlike the MG inoculation, the changes due to RR fruit inoculation correlated poorly with ripening (adjusted *r*^2^ = 0.02, **Supplemental Figure 2**). Accordingly, enrichment in positive log2 fold changes of pectin-related categories as a result of RR inoculation was weaker in contrast to MG fruit inoculation and ripening. Notably, RR inoculation did result in enrichment of negative log_2_ fold changes in xyloglucan-related categories, similar to ripening (**Supplemental Figure 2**).

### *B. cinerea* requires two pectin degrading enzymes to promote fruit susceptibility in unripe fruit

Transcriptomic and glycomic analyses indicated that pectin degradation is triggered by *B. cinerea* infections of MG fruit as well as fruit ripening, and that expression of a diversity of host CWDEs might be responsible. However, *B. cinerea* is known to employ its own cell wall degrading enzymes during infection to facilitate host tissue breakdown. Using the same samples from this study, we previously analyzed the *B. cinerea* transcriptome during infections of MG and RR tomato fruit (1 dpi and 3 dpi) and demonstrated that *B. cinerea* expression of pectin-degrading enzymes, particularly PGs (GH28 family), PL/PELs (PL1, PL3 families), and PMEs (CE8 family), is especially prominent during infections of MG fruit (Petrasch et al., 2019b). In **Supplemental Table S3**, we provide a list of differentially expressed fungal genes encoding key pectin degrading enzyme families based on the fungal RNAseq data. To pinpoint the individual genes from these families whose expression is prominent after MG inoculation and can contribute to the cell wall breakdown, we measured the expression of genes known to encode *B. cinerea* pectin-degrading enzymes by qPCR in fruit and vegetative tissues at 1 and 3 dpi (**Supplemental Figure 3**).

Four *B. cinerea* genes stood out as having high relative gene expression in MG fruit: *Bcpg*1, *Bcpg*2, *Bcpme*1, and *Bcpme*2. Bcpg1 and Bcpme1 are known virulence factors (ten Have et al., 1998; Valette-Collet et al., 2003), while Bcpg2 and Bcpme2 are comparatively understudied. To test the importance of Bcpg1, Bcpg2, Bcpme1, and Bcpme2 activity during MG inoculation, we utilized two previously reported double mutant *B. cinerea* lines *ΔBcpme*1*ΔBcpme*2 and *ΔBcpg*1*ΔBcpme*1, and the newly generated *ΔBcpg*1*ΔBcpg*2. We studied the double mutants instead of single ones because it is known that these CWDEs work interdependently to degrade pectin and some may present functional redundancy (Kars and van Kan, 2007). We evaluated their virulence in MG fruit by measuring disease incidence and severity each day from 3 to 6 dpi. All mutant strains except for *ΔBcpg*1 *ΔBcpg*2 were equally virulent as the wild-type strain on MG fruit (**Fig. 4**). *ΔBcpg*1*ΔBcpg*2 was completely avirulent on MG fruit, suggesting that the double knockout of these two PG genes was sufficient to prevent colonization on MG fruit. The importance of the cell wall integrity in unripe fruit to limit fungal infection is further supported by the fact that *ΔBcpg*1*ΔBcpg*2, as well as the other mutants, are completely capable of infecting RR fruit, which have partially disassembled cell walls (**Fig. 4**).

**Figure 4:**
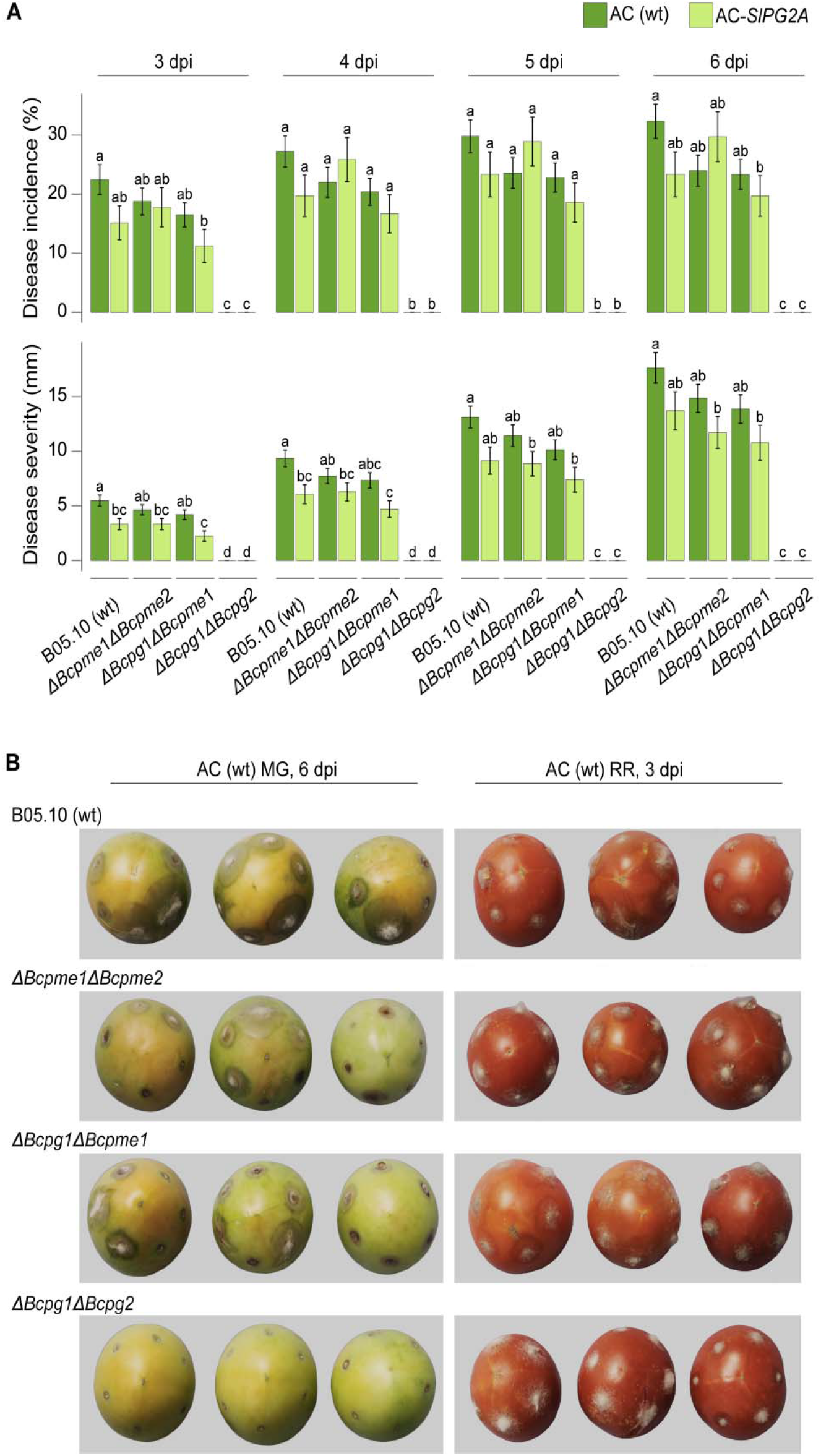
Disease incidence and severity of *B. cinerea* double mutants in unripe fruit. (**A**) Measurements of disease incidence and severity in wild-type (B05.10) and double mutants on wild-type (AC) and *AC-SlPG2A* MG fruit from 3 to 6 dpi (n = 55 - 161). Letters indicate statistical differences (*P* < 0.05) between *B. cinerea* and tomato genotypes at each dpi as calculated by ANOVA and Tukey’s HSD test. (**B**) Representative photos of inoculated wild-type MG fruit at 6 dpi (left) and wild-type RR fruit at 3 dpi (right). The background of the photographs was removed but the fruit images were not altered in any way and they were all processed equally. *ΔBcpg = B. cinerea* polygalacturonase mutant, *ΔBcpme* = *B. cinerea* pectin methylesterase mutant.

In addition to wild-type (AC) fruit, we also tested AC-*SlPG2A*, a tomato line with suppressed expression of the main ripening-associated PG (Smith 1990) and the highest expressed CWDE in RR fruit (average normalized read count = 16,328.9), in order to evaluate how the loss of this host enzyme would impact pathogen establishment and growth in MG fruit. Silencing of *SlPG2A* on its own does not improve resistance to *B. cinerea* in RR fruit (Cantu et al., 2008; Silva et al., 2021), but the importance of inducing pectin degradation during MG fruit infections may reveal a greater impact for this enzyme. Except for *ΔBcpg*1*ΔBcpg*2, which was completely avirulent on AC-*SlPG2A* fruit, all *B. cinerea* strains showed both reduced disease incidence and disease severity on AC*-SlPG2A* fruit compared to wild-type fruit (**Fig. 4**). This underscores that cell wall breakdown in inoculated MG fruit is the result of both host and pathogen CWDE activity and further highlights the importance of inducing host ripening processes during infection of unripe fruit.

### Induction of ripening by *B. cinerea* in MG fruit is dependent on Bcpg1 and Bcpg2

If the *ΔBcpg*1*ΔBcpg2* mutant is completely avirulent on MG fruit, we expected that inoculation of MG fruit with this strain would not accelerate ripening to the same degree that the wild-type strain (B05.10) did. To test this hypothesis, we performed additional assays using the same phenotypic analyses of ripening progression as before using *ΔBcpg*1*ΔBcpg*2-inoculated MG fruit and compared these to both *B05.10*-inoculated and mock-inoculated MG fruit (**Fig. 5**). All three measurements indicate that although *ΔBcpg*1*ΔBcpg*2 inoculation can accelerate ripening, it is not to the same degree as the B05.10 strain. Measurements in *ΔBcpg*1*ΔBcpg*2-inoculated fruit were found to be closer to mock-inoculated than B05.10-inoculated fruit. Furthermore, *ΔBcpme*1*ΔBcpme*2 and *ΔBcpg*1*ΔBcpme*1 had a similar impact on color progression compared to B05.10 (**Supplemental Figure 4**), suggesting that the loss of virulence in the *ΔBcpg*1*ΔBcpg*2 mutant is responsible for the weakened ability to promote ripening.

**Figure 5:**
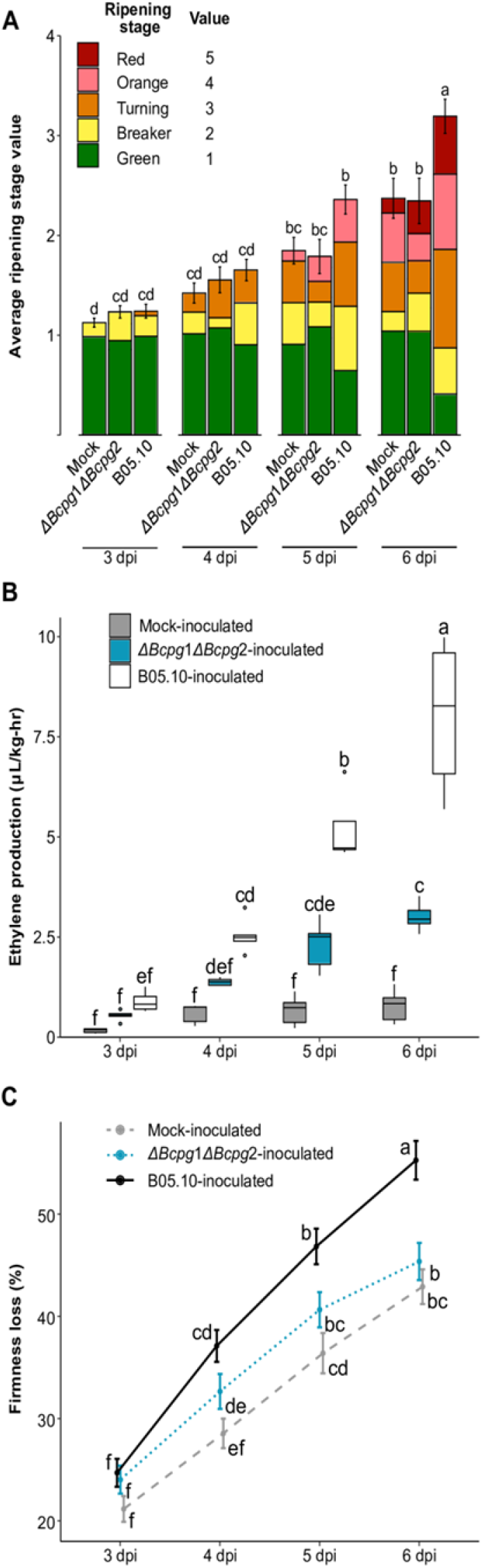
Ripening progression of unripe fruit inoculated with the *B. cinerea ΔBcpg*1*ΔBcpg*2 double mutant. **(A)** Average ripening stage value as assessed by color in mock-inoculated (M), B05.10-inoculated, and *ΔBcpg*1*ΔBcpg*2-inoculated MG fruit (n = 43 - 55) from 3 to 6 dpi. Colored blocks within each column represent the proportion of fruit at that respective stage. (**B**) Production of ethylene in M, B05.10-inoculated, and *ΔBcpg*1*ΔBcpg*2-inoculated MG fruit (n = 35 - 37). (**C**) Firmness loss in M, B05.10-inoculated, and *ΔBcpg*1*ΔBcpg*2-inoculated MG fruit measured as a percentage of initial firmness at 0 dpi (n = 70 - 216). Letters in **B-D** indicate the statistical differences (*P* < 0.05) between each treatment across all dpi as calculated by ANOVA and Tukey’s HSD test.

The failure of the *ΔBcpg*1*ΔBcpg*2 mutant to substantially induce ripening by 6 dpi suggests that it loses critical virulence factors for establishment and survival in MG fruit. However, it was unclear whether this loss is fatal or detrimental to the pathogen, or if it can remain quiescent until the fruit ripens at its normal rate, and then cause disease. To identify the ultimate fate of *ΔBcpg*1*ΔBcpg*2 mutants on MG fruit, we assessed disease incidence and measured fungal biomass in *ΔBcpg*1*ΔBcpg*2-inoculated fruit up to 20 dpi, well after the fruit reached the RR stage through normal ripening. These measurements revealed that *ΔBcpg*1*ΔBcpg*2 never fully recovered from its failure to substantially accelerate ripening, reaching a biomass at 20 dpi approximately only twice as great as its biomass at 3 dpi (**Table 3**) and there was no lesion development in any of the inoculated tomatoes. In contrast, when RR fruit were inoculated with *ΔBcpg*1*ΔBcpg*2 directly, the pathogen grew rapidly, and biomass at 3 dpi was nearly 350 times greater than the biomass at 20 dpi from inoculated MG fruit left to ripen. As a comparison, MG fruit inoculated with the wild-type strain B05.10 showed a biomass of 857.1 (± 109.74) μg/g fresh weight at 3 dpi, nearly 39 times greater than *ΔBcpg*1*ΔBcpg*2 on MG fruit at this timepoint, indicating that *ΔBcpg*1*ΔBcpg*2 growth is truly inhibited early after inoculation. Altogether, these results suggest that induction of ripening in MG fruit is a critical survival and infection strategy of *B. cinerea*, and that Bcpg1 and Bcpg2 are necessary for this strategy.

**Table 3:** Fungal biomass of *ΔBcpg*1*ΔBcpg*2-inoculated unripe fruit. Letters indicate statistical differences (*P* < 0.05) between each dpi among *Δbcpg*1*Δbcpg*2-inoculated MG fruit as calculated by ANOVA and Tukey’s HSD test (n = 3). Average fungal biomass determined in *ΔBcpg*1*ΔBcpg*2-inoculated Red Ripe RR fruit at 3 dpi is given as a reference (n = 4).

| Stage at Inoculation | dpi | Biomass ($\mu\text{g/g}$ fresh weight) |
| --- | --- | --- |
| MG | 3 | $22.1 \pm 3.8$ (b) |
| | 6 | $29.4 \pm 14.6$ (ab) |
| | 10 | $50.6 \pm 12.2$ (a) |
| | 15 | $30.1 \pm 8.3$ (ab) |
| | 20 | $43.3 \pm 8.9$ (ab) |
| RR | 3 | $14,951.6 \pm 3,386.5$ |

## Discussion

### Acceleration of tomato fruit ripening as a fungal infection strategy

The enormous benefit of ripening to infection success presents the opportunity for the evolution of a pathogen infection strategy that actively accelerates this process in unripe fruit. Such a strategy would suggest manipulation of host gene expression, as has been demonstrated by *B. cinerea* in vegetative and fruit tissues. To promote senescence, *B. cinerea* infection induces expression of genes associated with programmed cell death in tomato leaves (Hoeberichts et al., 2003). *B. cinerea* also actively suppresses host defense genes in both tomato and Arabidopsis through the production of small RNAs (Weiberg et al., 2013). In *B. cinerea*-inoculated unripe tomato fruit, previous microarray experiments have revealed upregulation of a small selection of ripening genes, including the ethylene biosynthesis genes *SlACS2* and *SlACS4*, the CWDEs *SlPG2A* and *SlEXP1*, and several others (Cantu et al., 2009). Thus, induction of ripening processes by *B. cinerea* to promote susceptibility is plausible.

Physiological measurements of hallmarks of climacteric fruit ripening confirm that *B. cinerea* can accelerate ripening in unripe tomato fruit (**Fig. 1**). While unripe fruit are resistant up to 3 dpi, the ripening acceleration that occurs after this point coincides with the onset of disease symptoms as *B. cinerea* emerges from quiescence and into its necrotrophic phase on these increasingly susceptible fruit. Additionally, RNA data at 1 dpi and 3 dpi support the activation of both ethylene biosynthesis and cell wall degradation. Color progression may be partly explained by high baseline expression of *SlPSY1* in MG fruit at 3 dpi, together with more significant transcriptional activity at later time points (**Supplemental Figure 1**).

*B. cinerea* inoculation of MG fruit at 1 and 3 dpi does not accelerate the expression of all known ripening-related genes. This is supported by the fact that we did not detect the upregulation of *SlACS2* (*Solyc02g068490*; **Supplemental Table S2**), while *SlPG2A* and *SlPL* (**Fig. 2**) were only demonstrably different at later time points (**Supplemental Figure 1**). Additionally, *B. cinerea* inoculation did not result in upregulation of the ripening-promoting transcription factors *SlRIN* (*Solyc05g012020*), *SlNOR* (*Solyc10g006880*), *SlCNR* (*Solyc02g077920*), or *SlTAGL1* (*Solyc07g055920*; **Supplemental Table S2**). However, it is possible that these genes as well as other ripening-related processes are triggered by inoculation at a different time point than those evaluated, as the accelerated ethylene biosynthesis will inevitably activate most ripening processes.

Ethylene is the most important hormone involved in climacteric fruit ripening and plant defense against pathogens, though it can promote either resistance or disease depending on the pathosystem (van der Ent and Pieterse, 2012). At the onset of ripening, fruits transition from System 1 (autoinhibitory biosynthesis) to System 2 (positive feedback loop) of ethylene production, leading to a burst in ethylene (Liu et al., 2015). In unripe tomato fruit, System 1 may facilitate resistance to *B. cinerea* up to 3 dpi, while System 2 accelerates ripening and promotes disease (Blanco-Ulate et al., 2013). The rate of increased ethylene biosynthesis observed in *B. cinerea*-inoculated unripe fruit after 3 dpi suggests that *B. cinerea* pushes the fruit into System 2 prematurely (**Fig. 1**). This is supported by the upregulation of two System 2 genes, *SlACS2* and *SlACO4*, in response to *B. cinerea* inoculation of unripe fruit.

### *B. cinerea* hijacks the host cell wall degrading machinery to facilitate fruit colonization

*B. cinerea* inoculation of MG fruit also results in upregulation of 34 different host CWDEs, which together with enzymes secreted by *B. cinerea* are likely responsible for the accelerated rate of softening in MG fruit. In particular, the mass upregulation of multiple host PMEs may be critical, as these enzymes are thought to facilitate further degradation by other enzyme classes (Jolie et al., 2010). Silencing of host CWDEs, particularly *SlPL*, reduces susceptibility to *B. cinerea* in ripe fruit (Silva et al., 2021). Thus, *B. cinerea* relies on host CWDEs to promote infection. This is further supported here by the reduced virulence of wild-type and mutant *B. cinerea* strains in MG fruit of the *AC-SlPG2A* mutant, which has silenced expression of *PG2A*, an important CWDE with extremely high expression levels in ripening fruit (Silva et al., 2021).

The glycomics analysis revealed that changes in the cell wall structure as a result of MG inoculation were remarkably similar to those that occur during ripening, particularly among the pectin polysaccharides (**Fig. 3**). Increased binding signals of RG I and its subcomponents in several fractions in both MG inoculation and ripening indicate increased accessibility to these molecules, perhaps due to loosening of the network through PMEs and/or degradation of side chains by various enzymes. The lack of substantial hemicellulose remodeling because of MG inoculation underscores the importance of the pectin network in mediating protection against *B. cinerea*. Interestingly, though the changes in cell wall structure as a result of RR infection were less similar to those caused by ripening, there were some similarities on xyloglucan depolymerization (**Supplemental Figure 3**). This suggests that the hemicellulose network is a secondary target for degradation by *B. cinerea* that receives focus after most pectin has already been broken down. Our previous research has also shown that expression of xyloglucanases by *B. cinerea* as well as two other pathogens, *Rhizopus stolonifer* and *Fusarium acuminatum*, appears more prevalent in infections of RR fruit than MG fruit (Petrasch et al., 2019b).

### Successful growth of *B. cinerea* in unripe fruit is dependent on its ability to degrade pectin

The importance of pectin degradation in MG fruit infections is a critical narrative emerging from the *B. cinerea-tomato* pathosystem. High prevalence of pectin-degrading enzymes, particularly polygalacturonases, in the expression profiles of *B. cinerea* on MG fruit was discovered previously (Petrasch et al., 2019b). Multiple pectin-degrading enzymes are expressed at high levels in MG fruit, and expression of these genes is greater on MG fruit compared to RR fruit and leaves (**Supplemental Figure 3**). While Bcpg1 and Bcpme1 are known to be important virulence factors, only the combined elimination of Bcpg1 and Bcpg2 resulted in complete avirulence on MG fruit despite the fact that *ΔBcpg*1*ΔBcpg*2 mutants are capable of causing disease on ripe fruit (**Fig. 4**; **Table 2**). Critically, infections of MG fruit by *ΔBcpg*1*ΔBcpg*2 do not appear to accelerate ripening processes to nearly the same degree as wild-type *B. cinerea* (**Fig. 5**).

These findings lead to several hypothetical contributing factors for pathogen-accelerated fruit ripening. First, early establishment of a quiescent infection in MG fruit is dependent on Bcpg1 and Bcpg2. Once established, *B. cinerea* may actively accelerate ripening through the secretion of unknown virulence factors. Additionally, in the absence of a quiescent infection, the host does not detect the pathogen and the lack of a response (e.g., ethylene-mediated signaling pathways), results in no trigger for early ripening. Bcpg1 and Bcpg2 may be required for substantial accumulation of pectin-derived oligosaccharides (PDOs) from the breakdown of pectin, as has been previously indicated (An et al., 2005). Although PDOs may function as signaling molecules during plant defense (Ferrari et al., 2013), they likely also act as triggers of ripening through elicitation of ethylene biosynthesis (Melotto et al., 1994).

These physiological, gene expression, and glycomics data all demonstrate that *B. cinerea* can induce ripening in MG fruit and uses this capacity to emerge from quiescence and cause disease. The induction of ripening appears dependent on the ability of *B. cinerea* to establish in unripe tissues even before causing lesion development. The pectin degrading enzymes Bcpg1 and Bcpg2 are key virulence factors as they are both critical for successful infection of MG fruit.

This research expands the understanding of pectin degradation in the *B. cinerea-tomato* pathosystem by suggesting that its importance goes beyond simply opening cell walls for colonization but also might trigger a cascade of ripening activities that cause the host to make itself more susceptible. This new dynamic may further guide identification of possible ripening-promoting virulence factors in *B. cinerea* and perhaps other postharvest fruit pathogens, and will ultimately improve our understanding of ripening-related susceptibility.

## Materials and Methods

### Biological material

Tomato (*Solanum lycopersicum*) cv. Ailsa Craig (AC) was obtained from the Tomato Genetics Research Center (UC Davis, USA). The *SlPG2A* antisense line in the AC background (*AC-SlPG2A*) was provided by D. Grierson (University of Nottingham, UK; Smith et al., 1990). Tomato plants were grown under typical field conditions during the summers of 2010, 2013, 2020, and 2021 in Davis, California. Tomato fruit from AC and AC-*SlPG2A* plants were tagged at 3 days post-anthesis (dpa) and harvested at 31 dpa for mature green (MG) and at 42 dpa for red ripe (RR) stages. The ripening stages were further confirmed by color, size, and texture of the fruit as in Adaskaveg et al. (2021).

The *ΔBcpg*1*ΔBcpg*2 mutant was produced by transforming the *ΔBcpg*1 mutant (ten Have et al., 1998) with a *Bcpg2* gene replacement construct containing a nourseothricin resistance (NAT) cassette. The sequences and details of the primers used for transformation are available in **Supplemental Table S4**. The *B. cinerea* B05.10 strain, the *ΔBcpg*1*ΔBcpg*2 mutant, as well as the *ΔBcpg*1*ΔBcpme*1 and *ΔBcpme*1*ΔBcpme*2 isogenic mutant strains (Kars et al., 2005a; Kars et al., 2005b) were grown on 1% potato dextrose agar as described in Petrasch et al. (2019b).

### *B. cinerea* inoculation

Tomato fruit were disinfected and inoculated as in Cantu et al. (2008). Fruit were wounded at six sites (depth of 2 mm and diameter of 1 mm) and inoculated with 10 μL of 5×10^5^ conidia/mL suspension of the wild-type strain (B05.10) or each of the mutants. Tomato fruit used as mock-inoculated material had 10 μL of sterile water placed on the wounds. Healthy fruit were not wounded or inoculated. MG and RR tomato fruit (i.e., *B. cinerea*-inoculated, mock-inoculated or healthy) were incubated at 20 °C in high humidity for different periods of times depending on the analyses.

Tomato fruit used for fungal biomass measurements, for transcriptomics, qPCR, and glycomics were deseeded, frozen and ground to fine powder in liquid nitrogen. Three to six biological replicates were produced per treatment and ripening stage; each consisted of independent pools of 8-12 tomato fruit.

### Assessments of ripening progression

For color progression, photos were taken of all fruit, and each individual fruit was visually categorized each day into one of five color groups each with a corresponding ripening stage value: mature green (1), breaker (2), orange (3), pink (4), and red ripe (5). These fruit were also assessed for the presence of fungal disease symptoms (e.g., water-soaked lesions). For ethylene, fruit were weighed each day and pooled into five airtight sterile containers as in Adaskaveg et al. (2021) and analyzed in a CG-8A gas chromatograph (Shimadzu Scientific Instruments, Kyoto, Japan). Ethylene production was calculated from the peak height, fruit mass, and incubation time. For firmness, fruit were assessed each day, as well as at 0 days, on the TA.XT2i Texture Analyzer (Texture Technologies, United States) using a TA-11 acrylic compression probe, a trigger force of 0.035 kg, and a test speed of 2.00 mm/sec. Firmness loss was calculated as the percentage of firmness at 0 days for each individual. Significant differences in physiological parameters between treatments were determined with analysis of variance (ANOVA) followed by post hoc testing (Tukey’s honestly significant difference, HSD) using R (R Foundation for Statistical Computing, Austria).

### RNA isolation and sequencing

Two grams of frozen ground fruit tissue (pericarp and epidermis) were used for RNA extraction, as described in Blanco-Ulate *et al*., 2013. RNA concentration and purity were measured using the NanoDrop 2000c Spectrophotometer (Thermo Fisher Scientific, USA). RNA integrity was checked by agarose gel electrophoresis. Eighteen cDNA libraries were prepared using the Illumina TruSeq RNA Sample preparation Kit v.2 according to the low-throughput protocol (Illumina, USA). Each library corresponded to three biological replicates of wild-type tomato fruit at MG and RR stages 3 days after treatment. The cDNA libraries were barcoded individually and analyzed for quantity and quality with the High Sensitivity DNA Analysis Kit in the Agilent 2100 Bioanalyzer (Agilent, USA). cDNA libraries were pooled in equal amounts for sequencing (single end, 50 bp) at the Expression Analysis Core Facility (UC Davis, USA) in an Illumina HiSeq 2000 sequencer.

### RNAseq data processing and functional analysis

Raw sequencing reads were trimmed for quality and adapter sequences using Trimmomatic v0.33 (Bolger et al., 2014) with the same parameters as reported in Silva et al., 2021. Trimmed reads were mapped using Bowtie2 (Langmead and Salzberg, 2012) to a combined transcriptome of tomato (SL4.0 release; http://solgenomics.net) and *B. cinerea* (http://fungi.ensembl.org/Botrytis_cinerea/Info/Index). Count matrices were made from the Bowtie2 results using sam2counts.py v0.91 (https://github.com/vsbuffalo/sam2counts/). Only reads that mapped to the tomato transcriptome were used in the following analyses. The Bioconductor package DESeq2 (Love et al., 2014) was used to normalize raw read counts and to determine differential expression (*P_adj_* < 0.05) among treatments. Differential expression results for 1 dpi data were obtained directly from Silva et al. (2021, GSE148217). For ripening gene expression, raw sequencing reads were downloaded from the fruitENCODE website (http://www.epigenome.cuhk.edu.hk/encode.html) and processed as above, with the exception that these reads were mapped only to the tomato transcriptome. Gene annotations for tomato were taken from Silva et al. (2021). All functional enrichments were performed using Fisher’s test with resulting *P*-values adjusted following Benjamini and Hochberg (1995).

### Quantitative PCR (qPCR)

cDNA was prepared from the isolated RNA using M-MLV Reverse Transcriptase (Promega, USA) as in Petrasch et al. (2019b). The tomato *UBIQUITIN LIKe-1* (*SlUbq-like1*, *Solyc12g04474*) and the *B. cinerea RIBOSOMAL PROTEIN-LIKE5* (*BcRPL5, Bcin14g04230*) were used as reference genes for tomato and *B. cinerea*, respectively, and processed in parallel with the genes of interest. Primer efficiencies were confirmed to be above 90% as in Petrasch et al. (2019b). Specificity of the primers was checked by analyzing dissociation curves ranging from 60 °C to 95 °C. qPCR primer sequences can be found in **Supplemental Table S4**. Transcript levels for all genes were linearized using the formula 2^(*REFERENCE* CT – *TARGET* CT)^. Data presented are for 3-6 biological replicates. Differences in relative expression levels were assessed by ANOVA followed by Tukey’s HSD using R. Sixteen tomato genes encoding different tomato CWDEs were selected for qRT-PCR validation of the RNASeq data at 3 dpi (**Supplemental Table S5**). A strong correlation (*r* = 0.88) was obtained between the log_2_ fold change values from the RNASeq data and the qPCR data.

### Cell wall extraction and fractionation

Total cell walls were prepared from combined fruit pericarp and epidermis (15 g) as described by Vicente et al. (2007), with the following modifications: samples were boiled in 100% ethanol for 45 min, and the insoluble material was filtered through glass microfiber filters (Ahlstrom, Finland) rather than Miracloth. Three preparations of extracted walls were obtained per experimental class; each extraction was from an independent pool of fruit (6-10 fruit) from three different harvests. Sequential chemical extractions of the total cell wall material (alcohol insoluble residue, AIR) were performed as specified in Vicente et al. (2007) to obtain WSF (water-soluble fraction), CSF (CDTA-soluble fraction), NSF (Na_2_CO_3_-soluble fraction) and KSF (24% KOH-soluble fraction). All the extractions were done at room temperature and the 4% KOH-soluble fraction was omitted. There were six replications per sample class.

### Glycomic analysis of cell wall fractions

Total sugar content of the cell wall fractions was calculated by adding the content of uronic acids and of neutral sugars present in each of the samples. The uronic acid content was measured according to Blumenkrantz and Asboe-Hansen (1973), and neutral sugar content was determined by the anthrone method (Yemm and Willis, 1954). Measurements for each fraction were done in triplicate using a Synergy H1 Hybrid Multi-Mode Microplate Reader (Biotek, USA). All cell wall fractions were diluted to the same total sugar concentration for the glycomic experiments. Glycome profiling of the cell wall fractions was performed by high-throughput ELISAs with a toolkit of plant cell wall glycan-directed monoclonal antibodies (Pattathil et al., 2010) as described by Zhu et al. (2010). Categorization of antibodies was retrieved from Pattathil et al. (2010), Dallabernardina et al. (2017), and Ruprecht et al. (2017). Antibodies with a maximum binding signal less than 0.1 across all fractions and treatments were filtered from further analyses. For the scatterplot analysis of treatments, a linear regression model was fitted to the data and was tested for statistical significance (*P_adj_* < 0.05) in R. Enrichments were performed using Fisher’s test with resulting *P*-values adjusted following Benjamini and Hochberg (1995).

### Disease development assays

Wild-type or *AC-SlPG2A* MG tomato fruit inoculated with the B05.10 strain or one of the mutant strains (*ΔBcpg*1*ΔBcpg*2, *ΔBcpg*1*ΔBcpme*1 and *ΔBcpme*1*ΔBcpme*2) were assessed for disease symptoms starting at 3 dpi. Disease development was recorded as disease incidence (percentage of inoculation sites showing symptoms) and disease severity (diameter of the soft rot lesions). These susceptibility evaluations were repeated over the course of eight separate harvest dates using 10-15 fruit per experimental treatment. Differences in disease incidence and severity between tomato genotypes and *B. cinerea* strains at each dpi were assessed by ANOVA followed by Tukey’s HSD using R.

*B. cinerea* biomass was quantified using the QuickStix Kit (EnviroLogix, USA), which utilizes the monoclonal antibody BC12.CA4 (Meyer et al., 2000) as described by Blanco-Ulate et al. (2015). Three to six biological replicates of the distinct *B. cinerea*-inoculated tissues were measured. One gram of tissue (pericarp and epidermis) from each biological replication was suspended in the kit buffer, 2:1 m/v for samples without obvious symptoms of fungal infection, 1:5 m/v for MG samples and 1:120 m/v for RR samples. The intensity of the monoclonal antibody reaction was determined using the QuickStix Reader (EnviroLogix, USA) and converted into fungal biomass (μg g^-1^ fresh weight of fruit extracts).

## Accession Numbers

The novel datasets for this study have been deposited in the Gene Expression Omnibus (GEO) database under the accession GSE183836.

## Acknowledgements

We want to recognize Dr. John M. Labavitch (dec.), who was instrumental in providing ideas and inspiration for this manuscript and other fruit cell wall research. We want to thank Estefanía Vincenti for her assistance in performing fungal inoculations and genotyping the double mutants. We also acknowledge Elijah Lerner for performing inoculations and measurements of firmness and ethylene.

## Supplemental Material

**Supplemental Figure 1: qPCR-based expression of selected tomato ripening-associated genes after inoculation with *B. cinerea*.** Names of each gene are given above each graph. Asterisks indicate statistical differences (*, *P* < 0.05; **, *P* < 0.01; ***, *P* < 0.001) between mock-inoculated and *B. cinerea*-inoculated mature green fruit across 3 to 6 days post-inoculation (dpi), as calculated by t-test (n = 3-8). (A) Tomato carotenoid biosynthesis genes: phytoene synthase (*SlPSY1*) and zeta-carotene desaturase (*SlZDS*). (B) Tomato ethylene biosynthesis genes: 1-aminocyclopropane-1-carboxylic acid oxidases (*SlACO1, SlACO3*) and 1-aminocyclopropane-1-carboxylic acid synthase (*SlACS4*). (C) Tomato cell wall degrading enzymes: polygalacturonase (*SlPG2a*), pectate lyase (*SlPL*), and a pectin methylesterase (SlPMEU1).

**Supplemental Figure 2: Glycomics profiling of *B. cinerea*-inoculated RR fruit at 3 dpi.** (**A**) Heatmaps of mAB binding strength in healthy MG, healthy RR, and *B. cinerea*-inoculated RR fruit for polysaccharide-binding antibodies. mAB codes are given to the right of each heatmap row, with the recognized classes of cell wall polysaccharides indicated by colored shapes according to the given key. mAB listed in boldtype are those included in the scatterplot in panel **B**. Arrows within heatmap tiles indicate statistically significant (*P_adj_* < 0.05) increasing or decreasing antibody strength when compared via t-test to values in healthy RR fruit. (**B**) Scatterplot and linear regression model of log_2_ fold change (log_2_FC) values of mAB signals in the ripening (healthy RR / healthy MG) and RR infection (*B. cinerea*-inoculated RR / healthy RR) comparisons. (**C**) Enrichment of polysaccharide classes with statistically significant positive or negative log2FCs in each cell wall fraction for the ripening and RR infection comparisons. Numbers within each tile indicate the number of mABs with a statistically significant log_2_FC in that respective fraction and polysaccharide class. WSF = water-soluble fraction, CSF = CDTA-soluble fraction, NSF = Na_2_CO_3_-soluble fraction, KSF = KOH-soluble fraction.

**Supplemental Figure 3: qPCR-based expression of selected cell wall degrading enzymes expressed by *B. cinerea* during tomato infections.** Names of each gene are given to the right of each graph. Letters indicate statistical differences (*P* < 0.05) between tissues across both 1 and 3 days post-inoculation as calculated by ANOVA and Tukey’s HSD test (n = 4 - 6). (**A**) *B. cinerea* polygacturonase *(BcPG)* and rhamnogalacturonase (*BcRG*) genes. (**B**) *B. cinerea* pectin methylesterase (*BcPME*) genes. (**C**) *Botrytis cinerea* pectin lyase (*BcPL*) and pectate lyase (*BcPEL*) genes. MG = Mature Green, RR = Red Ripe, dpi = days post-inoculation.

**Supplemental Figure 4: Color progression in *B. cinerea* mutant-inoculated MG fruit.** Average ripening stage value as assessed by color in B05.10-, *Δbcpg1Δbcpg2-, Δbcpg1bcpme1*-, and *Δbcpme1bcpme2*-inoculated MG fruit (n = 55 - 128). Colored blocks within each column represent the proportion of fruit at that respective stage. Letters indicate statistical differences (*P* < 0.05) between each treatment across all dpi as calculated by ANOVA and Tukey’s HSD test. **Supplemental Table S1.** Ripening stage assessments of mock-inoculated and *B. cinerea*-inoculated fruit each day from 3 to 6 dpi. *B. cinerea*-inoculated fruit are further divided into symptomatic (i.e., those exhibited water-soaked lesions) and asymptomatic fruit. Each count represents an individual tomato. Corresponding numerical values for each ripening stage value are provided in parentheses.

**Supplemental Table S2. Differential expression output from DESeq2 (Love et al., 2014) with functional annotations.** Comparisons are listed in the header of each section. Subheaders are values returned by the results function in DESeq2. baseMean = the mean of normalized counts of all samples for that gene; log2FoldChange = the logarithm (base 2) of the fold change between healthy and infected samples; lfcSE = the standard error of the log2FoldChange; stat = the Wald test statistic for each comparison; *P* = *P* value generated from the Wald test statistic; *P*adj = adjusted *P* value as calculated via the Benjamini & Holchberg method. AHRD = Automated Assignment of Human Readable Descriptions; KEGG = Kyoto Encyclopedia of Genes and Genomes. MG = Mature Green. N = Not differentially expressed in that comparison.

**Supplemental Table S3. Differential expression output of *B. cinerea* genes adapted from Petrasch et al., 2019b. Genes presented are those from CE8, GH28, PL1, and PL3 CAZY families with significant upregulation compared to in vitro samples for at least one comparison.** Comparisons are listed in the header of each section. Subheaders are values returned by the results function in DESeq2. baseMean = the mean of normalized counts of all samples for that gene; log2FoldChange = the logarithm (base 2) of the fold change between healthy and infected samples; *P*adj = adjusted *P* value as calculated via the Benjamini & Holchberg method.

**Supplemental Table S3. Differential expression output of *B. cinerea* genes adapted from Petrasch et al., 2019b. Genes presented are those from CE8, GH28, PL1, and PL3 CAZY families with significant upregulation compared to in vitro samples for at least one comparison.** Comparisons are listed in the header of each section. Subheaders are values returned by the results function in DESeq2. baseMean = the mean of normalized counts of all samples for that gene; log2FoldChange = the logarithm (base 2) of the fold change between healthy and infected samples; *P*adj = adjusted *P* value as calculated via the Benjamini & Holchberg method.

**Supplemental Table S4. Primers used for genotyping *B. cinerea* mutants and qPCR expression analyses of *B. cinerea* and tomato genes**

**Supplemental Table S5. Correlation between qPCR and RNASeq log2FoldChanges for 16 chosen tomato genes.** log2FoldChange = the logarithm (base 2) of the fold change between treatments; AHRD = Automated Assignment of Human Readable Descriptions; KEGG = Kyoto Encyclopedia of Genes and Genomes.

## Notes

**Funding Information:** This work was supported by start-up funds from the College of Agricultural and Environmental Sciences and the Department of Plant Sciences (UC Davis) to BB-U. Funding to CJS was partially provided by the Plant Sciences Graduate Student Researcher Award (UC Davis). This research was also supported by the National Science Foundation under grants IOS 0544504 and IOS 0957264 awarded to ALTP.

